# Aβ Fibrils Can Act as Aqueous Pores: a Molecular Dynamics Study

**DOI:** 10.1101/418137

**Authors:** S. Natesh, J. R. Sachleben, T. R Sosnick, K. F. Freed, S. C. Meredith, E. J. Haddadian

## Abstract

Aggregation of Aβ peptides is important in the etiology of Alzheimer’s Disease (AD), an increasingly prevalent neurodegenerative disease. We ran multiple ∼ 300 ns all-atom explicit solvent molecular dynamics (MD) simulations starting from three NMR-based structural models of Aβ(1-40 residues) fibrils having 2-fold (pdb code 2LMN) or 3-fold rotational symmetry (2LMP, and 2M4J). The 2M4J structure is based on an AD brain-seeded fibril whereas 2LMP and 2LMN represent two all-synthetic fibrils. Fibrils are constructed to contain either 6 or an infinite number of layers made using periodic images. The 6 layer fibrils partially unravel over the simulation time, mainly at their ends, while infinitely long fibrils do not. Once formed, the D23-K28 salt bridges are very stable and form within and between chains. Fibrils tend to retain (2LMN and 2LMP) or develop (2M4J) a “stagger” or register shift of β-strands along the fibril axis. The brain-seeded fibril rapidly develops gaps at the sides of the fibril, which allows bidirectional flow of water and ions from the bulk phase in and out the central longitudinal core of the fibril. Similar but less marked changes were also observed for the 2LMP fibrils. The residues defining the gaps largely coincide with those demonstrated to have relatively rapid Hydrogen-Deuterium exchange in solid state NMR studies. These observations suggest that Aβ(1-40 residues) fibrils may act as aqueous pores that might disrupt water and ion fluxes if inserted into a cell membrane.

## Introduction

Alzheimer’s Disease (AD) affects millions of people and is becoming more prevalent as the population ages (1-5). AD is believed to be mainly caused by aggregates of β-amyloid peptides and neurofibrillary tangles consisting of the intermediate filament protein, Tau (6-10). The end-points of aggregation are amyloid fibrils, though is it widely believed that soluble oligomers, including fibril precursors, are the more important neurotoxins (11-25).

Amyloid fibril formation is distinct from normal protein folding, where a single amino acid sequence generally folds to a single conformation; in contrast, amyloidogenesis yields polymorphous (heterogeneous) structures even from chemically pure proteins or peptides. Amyloids are defined by the presence of β-sheets, which commonly are parallel and in-register (26-29), but beyond this rudimentary structure motif, the three-dimensional arrangement of the β-sheets and intervening loops is polymorphic. Formation of diverse fibril structures depends on fibrillization conditions. This diversity appears to arise mainly at nucleation, since seeding fresh solutions of Aβ with pre-existing seeds leads to the formation of replicate fibrils, i.e., fibrils having the same structure as parental fibrils, by criteria such as transmission electron microscopy (TEM) and solid-state (SS) NMR (30-36).

The ability to form replicate fibrils from seeds was used to interrogate the structure of AD brain amyloid fibrils (35, 37). Amyloid harvested from the brains of two patients who had died with AD was used to seed the formation of replicate fibrils out of synthetic, isotopically labeled Aβ(1-40 residues). The replicate fibrils were then studied by biophysical methods, especially solid-state NMR spectroscopy. In each patient, replicate fibrils derived from amyloid from three different brain regions had a single structure, as judged from TEM and SS-NMR spectra, but each patient’s replicate fibril structure was distinct. In addition, a detailed structural model (2M4J) of the replicate fibrils from one patient differed from those of previously described, all-synthetic fibrils (10, 37).

Here, we compare MD simulations of the replicate fibrils based on brain-seeded nuclei (2M4J, (37) with two other Aβ fibril structural models (2LMN and 2LMP, (38,39) (Figure 1, Table 1). Fibrils 2LMN and 2LMP are all-synthetic polymorphs formed without seeding, starting with Aβ(1-40 residues) solutions, formed under either “agitated” (gently swirling) and “quiescent” fibrillization conditions, respectively. The 2LMN fibrils have a striated, linear appearance in TEM (9), and a two-fold axial symmetry. The 2LMP fibrils have a twisted ribbon appearance in TEM (9), and three-fold axial symmetry. Additional features of the two synthetic fibrils include the presence of a D23-K28 salt bridge in the 2LMN two-fold symmetric fibril, which is absent in the three-fold symmetrical 2LMP fibril, and present in the brain-seeded, 2M4J fibrils. In the 2LMN and 2LMP fibrils, the center of the N-terminal β-strand of peptide *i*, is coplanar with the center of the C-terminal β-strand of peptide *i*+*2*, thereby forming a β-strand stagger along the fibril axis. In the 2M4J fibrils, the N- and C-terminal β-strands of each molecule are initially co-planar.

**Table 1.** Characteristics of Structural Models of Aβ(1-40 residues) used for these studies.

| Name | PBD | Source | Symmetry | D23-K28 salt bridge | TEM appearance | Simulation |
| --- | --- | --- | --- | --- | --- | --- |
| Brain-seeded | 2M4J | Brain-seeded | 3-fold | Yes (6)* | Linear (untwisted) duplexes | Two infinites replicates (500 and 310 ns) and One finite (300 ns) |
| Synthetic 3-fold | 2LMP | All synthetic, “quiescent” | 3-fold | No (0) | Twisted ribbons | Three infinites replicates (300, 310 and 300 ns) One finite (300 ns) |
| Synthetic 2-fold | 2LMN | All synthetic, “agitated” | 2-fold | Yes (4) | Linear striated | Two infinites replicates (300 and 310 ns) two finite replicates (300 ns) |

\* The average number of initial D23-K28 salt bridges per stack in the NMR structures deposited in PDB. The initial 2M4J salt bridges are all intra-monomer and the 2LMN salt bridges are all inter-monomer.

**Figure 1.**
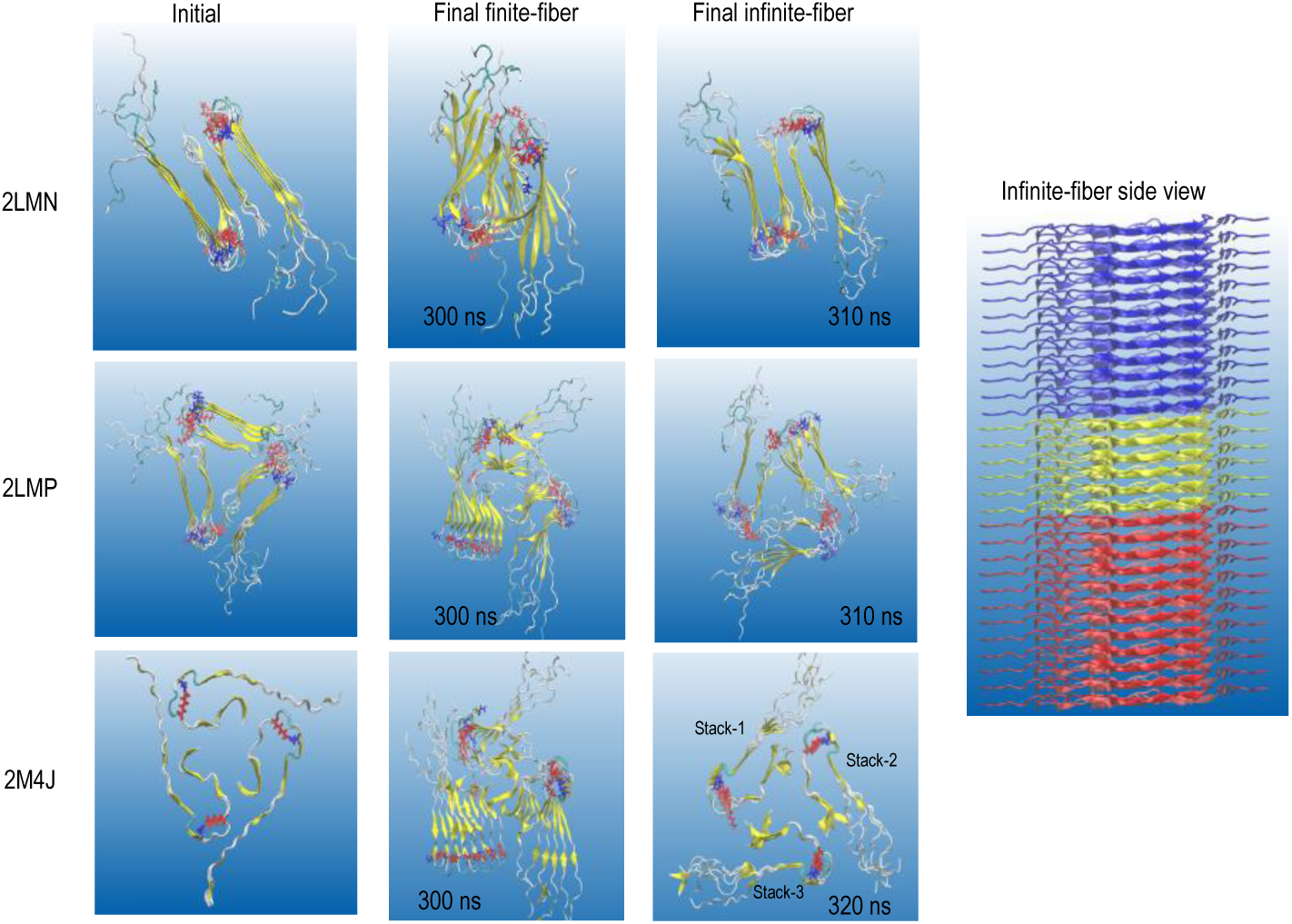
(A) Top view of the initial and final structures of three fibril models after the simulations. The three fibril types are 2LMN (two-fold symmetrical, all-synthetic), 2LMP (three-fold symmetrical, all-synthetic), and 2M4J (three-fold symmetrical, brain-seeded). The figure compares finite length fibrils containing six molecular layers, and infinitely long fibrils, constructed by replicating 6-layered structure along the *z*-axis using periodic images. Asp 23 (red) and Lys 28 (blue) form salt bridges in 2LMN and 2M4J, but not 2LMP. A side view of the 2M4J infinitely long fibril structure is also shown; the structure of the repeated unit is depicted in yellow, as are the two periodic images above and below in blue and red, respectively.

MD simulations have moved apace with structural studies of Aβ aggregates, mostly fibrils, and these computational studies recently have been thoroughly compared with experimental studies (40). However, many challenges still exist in this area, even for the relatively well understood fibrils. Most MD simulations have been performed using a number of layers that is orders of magnitude lower than those occurring in actual Aβ fibrils. Thus, the fibril ends might play a disproportionate role in such simulations, while events in the interior of the fibril could be overlooked.

To address this potential problem, we constructed infinitely long fibrils, using periodic images of 6-layer NMR fibril structures. We found that the finite fibrils undergo conformational rearrangements not observed in the infinite fibrils. The D23-K28 salt bridges in 2LMN and 2M4J remained mainly intact over the course of the simulations, while 2LMP fibrils acquired in average one such salt bridge during the course of simulations. The fibrils tend to retain (2LMN and 2LMP) or develop (2M4J) a stagger of β-strands in the long fibril axis (Z-axis). Most importantly, we observed a separation of the chains in the x-y plane for the three-fold symmetric brain-seeded fibril (2M4J), that led to penetration of solvent and ions in and out of the core of the fibril. This penetration was essentially absent in the other two fibrils. We discuss possible pathobiological consequences of these observations.

## Methods

### MD simulations

We performed all atom, explicit solvent (TIP3P) molecular dynamics simulations based on three structural models of amyloid Aβ(1-40 residues) fibrils (Table 1) using the CHARMM36 force field and NAMD 2.10 (41). These fibrils consist of 2 or 3 vertical “stacks” (in the z-axis) with each layer in the xy-plane containing either 2 (2LMN) or 3 (2LMP and 2M4J) molecules arranged with rotational symmetry about the z-axis. Finite length fibrils were comprised of 6 layers (12 or 18 chains total), unless otherwise noted. For the sake of consistency, we doubled the deposited 2M4J structure, which has only 3 layers, such that our simulations also were performed on 2M4J fibrils containing stacks of 6 layers. A total of 13 simulations were run as listed in Table 1. For each fibril model, the first conformer deposited in the Protein Data Bank was used. The N and C termini of Aβ monomers in all structures were protonated.

### Addition of missing residues

Residues 1-8 are missing in the structural models of the all-synthetic fibrils (2LMN and 2LMP), because they are not seen in SS-NMR spectra. These residues, however, were observed by SS-NMR for the brain-seeded fibrils (2M4J), and are shown in the structural models. For proper comparison, we constructed models of the all-synthetic fibrils containing these missing residues, using the program Modeller (42). In addition, we also constructed a 2LMP infinite length model without these residues (the N-terminus of Aβ monomers in this structure were protonated).

### Construction of infinite Aβ(1-40 residues) fibrils for simulations

Infinite length fibrils were constructed by replicating each 6-layered structure along the *z*-axis using periodic boundary conditions. The top and bottom Aβ molecules were allowed to interact directly with the adjacent images. In the 2LMN and 2LMP structures, the center of the N-terminal β-strand of peptide *i*, is coplanar with the center of the C-terminal β-strand of peptide *i*+*2*. In the 2M4J fibrils, the N- and C-terminal β-strands of each molecule are co-planar. This stagger in 2LMN and 2LMP led to atomic clashes when we attempted to form infinitely long fibrils using periodic images; this was especially obvious around the “bend” regions (amino acids 24-29). To separate the clashing loop regions, we rebuilt them with the loop modeling protocol implemented in the program Modeller (42). When necessary, we resolved inter-boundary side chain clashes by rotating them using the Molefacture plugin in VMD (43).

### Solvent conditions

The finite and infinite structures were solvated with a padding of 12.5 or 20 Å, respectively, between the protein and the boundary of the solvation box. The padding along the z-axis in infinite fibrils was set so that overlap among water molecules of periodic images was minimal. Any overlap disappeared during the system equilibration period. The 2LMP and 2M4J structures started with a water filled core. All systems were ionized to neutrality with NaCl. Harmonic constraints were placed on the protein backbone atoms and were removed gradually during the equilibration period.

### Preservation and formation of salt bridges

We tracked the preservation of salt bridges between the side chains of D23 and K28 in 2LMN and 2M4J fibrils, and also the possible formation of a salt bridge in 2LMP (all-synthetic, 3-fold symmetrical fibrils). The K28-D23 salt bridge was considered present if the distance between Nε of K28 was ≤ 3.7 Å from Oγ of D23 and if this criterion was fulfilled in at least 60% of the simulation time. Salt bridges can be intramolecular or intermolecular, or both. Hence, it is possible for a single D23 or K28 to be involved in more than one salt bridge.

As mentioned, loop regions of 2LMN were rebuilt to avoid steric clashes. The deposited 2LMN structure showed an average of four D23-K28 salt bridges per stack of six molecular layers. In the rebuilt loop regions of 2LMN we ensured that initial salt bridges were not distorted by maintaining a distance of ≤ 3.7 Å between Nε of K28 and Oγ of D23 during the loop remodeling.

### Simulation details

After constructing fibrils as described, all of the Aβ peptide backbone atoms were restrained using harmonic constraints, and the structures underwent energy minimization, using conjugate gradient, steepest descent for 20,000 steps. The infinite fibril systems were subsequently heated to 300K over 0.33 ns in the canonical (NVT) ensemble, followed by a gradual pressure application to atmospheric pressure and removal of the harmonic constraints from the loops and tails (residues 1-10 and 23-30) over 2.45 ns. The pressure applied to the system was increased in a gradual manner (from 0 bar to atmospheric pressure) to prevent structural deformation at the periodic boundaries and to preserve the spacing between periodic images; this procedure was necessary to simulate infinitely long fibrils. The finite fibril systems were heated to 300K for over 0.33 ns in the isobaric-isothermal (NPT) ensemble without a gradual pressure application. We simulated a 2LMN finite system with gradual pressure increase; the resulting model was very similar to the one with instant pressure application. After removing harmonic constraints on the rest of the protein (3.45 ns), the production simulations were performed in the NPT ensemble (1 bar and constant temperature *T* = 300 K). Simulations were regulated using Nosé-Hoover Langevin piston pressure control (44, 45) and Langevin damping dynamics. Bonded and short-range non-bonded interactions are calculated at every time step (2 fs). Electrostatic interactions were evaluated every 2 time steps using the particle mesh Ewald method (46). The cut-off distance for non-bonded interactions was 12 Å. The van der Waals interactions were smoothly truncated to 0 between 10-12 Å, while electrostatic interactions were shifted so that they had a value 0 at the cut-off distance. The pair list for non-bonded interaction was calculated every 10 time steps for those pairs separated by less than or equal to 13.5 Å. Distances of all bonds to hydrogen atoms were constrained by the RATTLE algorithm (47). The trajectories were analyzed with in-house scripts and the program VMD (43).

### Tracking water molecules

A central, hydrated core in the 2M4J structure is formed by residues 31-37. The core radius is computed using the program Hole (48, 49), which employs a Monte Carlo simulated annealing procedure to find the maximal radius of a sphere that can be ‘squeezed’ through the core at a given position along the core’s normal axis. Thus, Hole outputs a radius value for positions along the core’s normal (here, *z*) axis. We ran Hole at each frame of the trajectory and obtained radii for positions along the core’s normal axis for each frame. The reported radius was calculated as the average of radii corresponding to the middle three monomers that form the core for each frame.

We calculated the residency time for water molecules in the central core as the average time it takes for these molecules beginning in this region to exit. For a given reference frame, we selected water molecules in the core and started a frame counter for each molecule, terminating it upon the molecule’s exit. This procedure was run on multiple trajectory windows along the full trajectory and for windows of different sizes, from which an average and standard deviation were calculated.

As shown below, ions and water diffuse through both the edges and the central core of 2M4J (brain-seeded) fibrils. In order to compare water diffusion through and within the 2M4J fibrils to diffusion in bulk water, the mean square displacements of water molecules inside the 2M4J central core were calculated. For detailed description of the diffusion analysis, see Supporting Information.

We also tracked sodium ion permeation through the core. The coordinate space was partitioned into 5 regions, with the origin placed at the center of the core. All sodium ions were assigned labels that were updated every frame, corresponding to their location in the partitioned space. Based on how the labels changed throughout the trajectory, it could be determined whether a given sodium ion permeated the core.

### NMR Calculations

NMR and simulated structures were compared to the distance and backbone torsion angle constraints available through the protein databank for the structure 2M4J (filename 2m4j.mr). Comparisons were made by producing a histogram of distances for the atoms of a given constraint from the coordinate files. The NMR derived structure 2M4J consists of 20 structural models each of which contain 9 amyloid chains. For each given atom pair in a distance constraint, this produces 900 distances that were included in a histogram. For ambiguous constraints, all atom pairs were included in the histogram so that the total number of distances is 900 times the degeneracy of the constraint. The simulated structure included 18 amyloid chains, which produced 171 distances per atom pair of constraint. These distances were then averaged over the last 200 steps of simulation where it is assumed that the system has reached equilibrium. These histograms were then compared with the range of values given by the constraint file. If the histogram showed distances within this range, the structure is considered to be consistent with that constraint. Backbone torsion angle constraints were treated similarly. The constrained phi/psi angles were calculated from the 20 models of 2M4J, while for the simulations the phi/psi angles were averaged over the last 200 timesteps.

## Results

### Finite fibrils, but not infinite fibrils, unravel in simulations

For the three fibril forms (2LMN, 2LMP, and 2M4J), we compared the MD simulations of the infinite-length fibrils to fibrils comprised of 6-molecule layers of Aβ(1–40) peptides. Infinite-length fibrils were constructed by replicating each 6-layered structure along the *z* axis using periodic images.

All finite fibrils exhibited significant deviation from the starting structures, mainly an unraveling of the ends, whereas the infinite fibrils showed less of this deviation (**Figure 1**). In all cases, the infinite-length fibrils had lower final backbone RMSD values than their finite length counterparts (**Figure 2**). Among the finite fibrils, the 2M4J (brain-seeded) model had the maximum deviation from initial to final state. As shown in **Figure 2** for the central regions of the peptide (i.e., excluding the mobile terminals, residues 1-8 and 37-40), the RMSD of the finite fibrils increased by ∼ 8-11 Å, while the infinite fibrils, the increase was only 4-5 Å. The RMSD calculation including all residues also showed similar trends (supporting **Figure 1,** RMSD values for replicate simulations are also shown). In 2M4J fibrils (finite > infinite length, discussed below), the three stacks separated from each other. In the 2M4J and 2LMP fibrils, we also observed the loss of the initial 3-fold symmetry (**Figure 1**, finite >> infinite).

**Figure 2.**
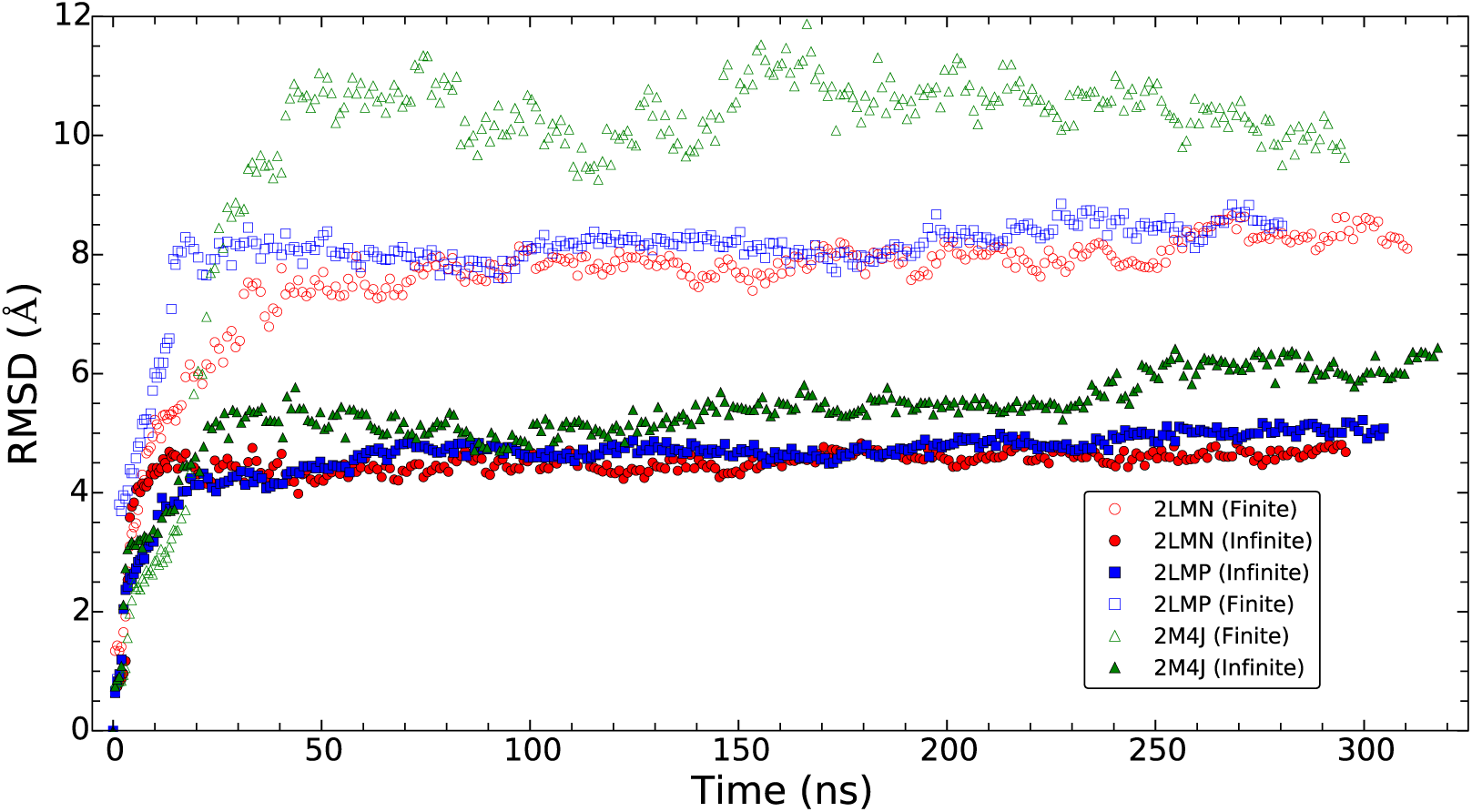
Backbone RMSD of fibrils. Residues 1-8 and 37-40, which are flexible and not observed in NMR studies of 2LMN (all-synthetic, two-fold symmetrical) and 2LMP (all-synthetic, three-fold symmetrical) (see, for example, 38, 39), are not included in the calculations. Finite fibrils are shown with open symbols, infinite fibrils with closed symbols. Red, blue and green correspond to 2LMN, 2LMP, and 2M4J, respectively. For clarity, every fiftieth point is plotted. RMSD values for replicate trajectories are shown in supporting Figure 1.

The initial jump in the RMSD values largely represents an equilibration period as the added initial restraints on the backbone atoms were gradually removed (the initial structure is a single configuration of an ensemble of structures; the MD modeling allows the structure to relax around this structure). The RMSD values for each fibril leveled off after the first 20-40 ns of the simulation time.

Similar findings were obtained using a quantification of the structural flexibility, the root-mean-square-fluctuation (RMSF) of Cα atoms (**Figure 2**). The RMSF values for the residues 9-36 for each fibril were computed for each monomer over the last 150 ns of the trajectory and then averaged over all monomers in a vertical stack. The average RMSF values in the finite fibrils are higher than in the infinite length fibrils, indicative of increased structural integrity for the infinite length models. In the finite fibrils, residues 22-30, corresponding to the loop regions of the Aβ peptides, had the largest fluctuations. We observed that the loop regions of the top and bottom layer monomers started to separate from the stack. As discussed below, the fraying of the ends of the finite length fibrils may have mechanistic implications for fibril disassociation. This distinction was not apparent in the infinite fibrils.

### Salt-bridges

A key feature of some fibrils, but not all, is a salt bridge between the side chains of D23 and K28 in the initial SS-NMR-based models. This feature was detected in fs-REDOR experiments of appropriately labeled peptides, and has been found in 2LMN (30, 38, 50) and 2M4J (37) fibrils, but not in 2LMP (39, Table 1). We tracked the salt bridges present initially in 2LMN and 2M4J and investigated whether salt bridges would develop in 2LMP fibrils. The number of D23-K28 salt bridges was nearly constant over the course of the simulation in all systems (**Supporting Figure 2A**). The number of initial salt brides in the 2LMN finite system decreased due to unraveling of the structure during the simulation. The 2LMP structures (both finite and infinite) had virtually no salt bridges initially and developed an average of only one salt bridge per stack, transiently, during the course of the simulations. In the 2M4J (brain-seeded) fibrils, initially all D23-K28 salt bridges were intramolecular. In the course of the simulations, however, in some cases the K28 side chain interacted with D23 side chains of two flanking molecules in a vertical stack, simultaneously forming intra- and intermolecular salt bridges (**Supporting Figure 2B**). Because K28 can interact simultaneously with D23 of two adjacent Aβ molecules, the infinite 2M4J fibril had more salt bridges than the other fibril types and acquired 3 additional salt bridges per stack during the course of the simulation. The significant deformation of the finite length 2M4J fibril during simulations, did not result in the reduction of its original number of D23-K28 salt-bridges. The number of D23-K28 salt-bridges in the infinite fibril trajectory replicates are almost the same in each of the fibril types (Supporting Figure 2C).

### Missing residues

Residues 1-8 are missing in the structural models of the all-synthetic fibrils because they are seen either poorly or not at all in the SS-NMR experiments. They are also not considered in most of computational simulations of Aβ peptides. These residues, however, were observed by SS-NMR of brain-seeded fibrils and are shown in the structural model derived from NMR and other experimental data (37). **Supporting Figure 3A** shows that deletion of these residues increased the structural deformation of the infinite-length fibril. The structural flexibility was also increased in one of the 32-residue fibril stacks (larger RMSF values in stack-1, **supporting Figure 3B)**. This region of the Aβ(1–40) contains a group of five charged residues that formed several transient salt bridges in the course of simulations helping to stabilize the 2LMP structure.

**Figure 3.**
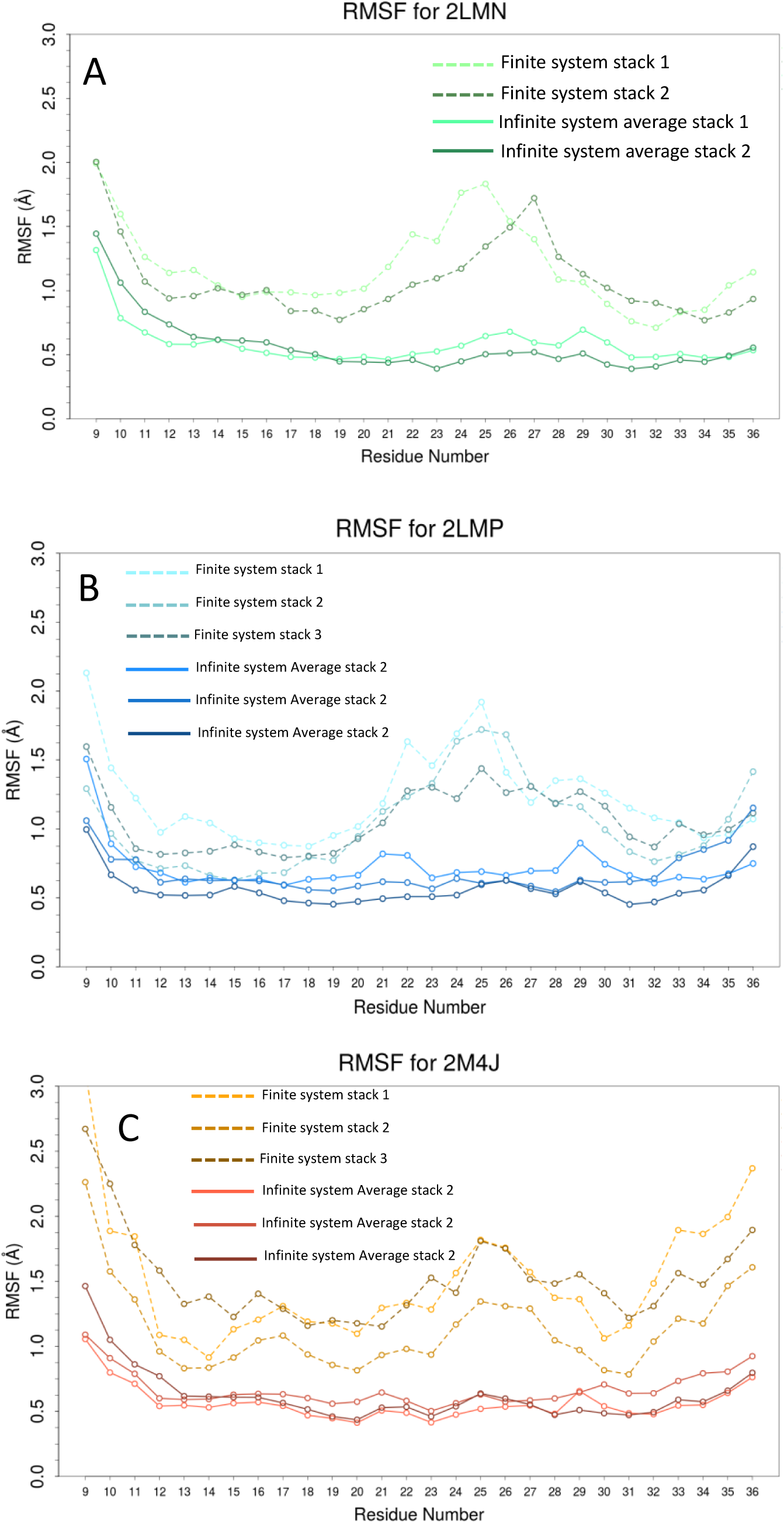
Cα-RMSF of fibrils (as in **Figure 2**, flexible residues 1-8 and 37-40 are excluded). Dashed lines are finite length fibrils solid lines are infinite fibrils. A: 2LMN (all-synthetic, two-fold symmetrical). B: 2LMP (all-synthetic, three-fold symmetrical). C: 2M4J (brain-seeded, three-fold symmetrical). Average RMSF values for residues in each stack in different replicates are shown.

### Alterations in the stagger

In the 2LMN and 2LMP structures, the center of the N-terminal β- strand of each peptide *i* is coplanar with the center of C-terminal β-strand of peptide *i*+*2*, i.e., there is a “stagger” in the fibril structure. A consequence of this stagger is that the fibril ends contain solvent-exposed hydrophobic residues, as well as unsatisfied hydrogen bonding sites. In contrast to 2LMN and 2LMP, in 2M4J fibrils the N- and C-terminal β-strands of each molecule are initially co-planar. During the simulation, the molecular stacks of 2M4J, especially in the finite-length fibrils, tended to develop a stagger reminiscent of those in the all-synthetic fibrils (**Supporting Figure 4**). This also occurred, albeit at a slower rate and in only one of the three stacks, in both replicate simulations of the infinite-length 2M4J fibril (See Stack 3, arrow, **Supporting Figure 4**).

### Movement of water molecules in and through fibrils

The β-sheet regions of many amyloids, notably of Aβ, are comprised mainly of hydrophobic amino acids. Furthermore, some interfaces between β-sheet regions of amyloids are thought to be nearly anhydrous (51-54). This was also the case for the β-sheet regions of the fibrils examined in our simulations. Water molecules were excluded from the regions occupied by side chains between β-sheet lamina. On the other hand, water molecules were initially present within the center of the 2M4J, and 2LMP models (Figure 4A, both finite- and infinite-length). The number of water molecules in the central core of the 2M4J fibrils (both the finite and infinite length) significantly increased because of the gaps that opened at the sides of the fibrils between the N-terminal region of one peptide, and the bend region of an adjacent peptide (**Figure 4A**). There was greater disruption in the finite-length 2M4J fibrils, where the stacks became more disordered (**Figure 1**). The gap opening was small for both the finite and infinite 2LMP models.

**Figure 4.**
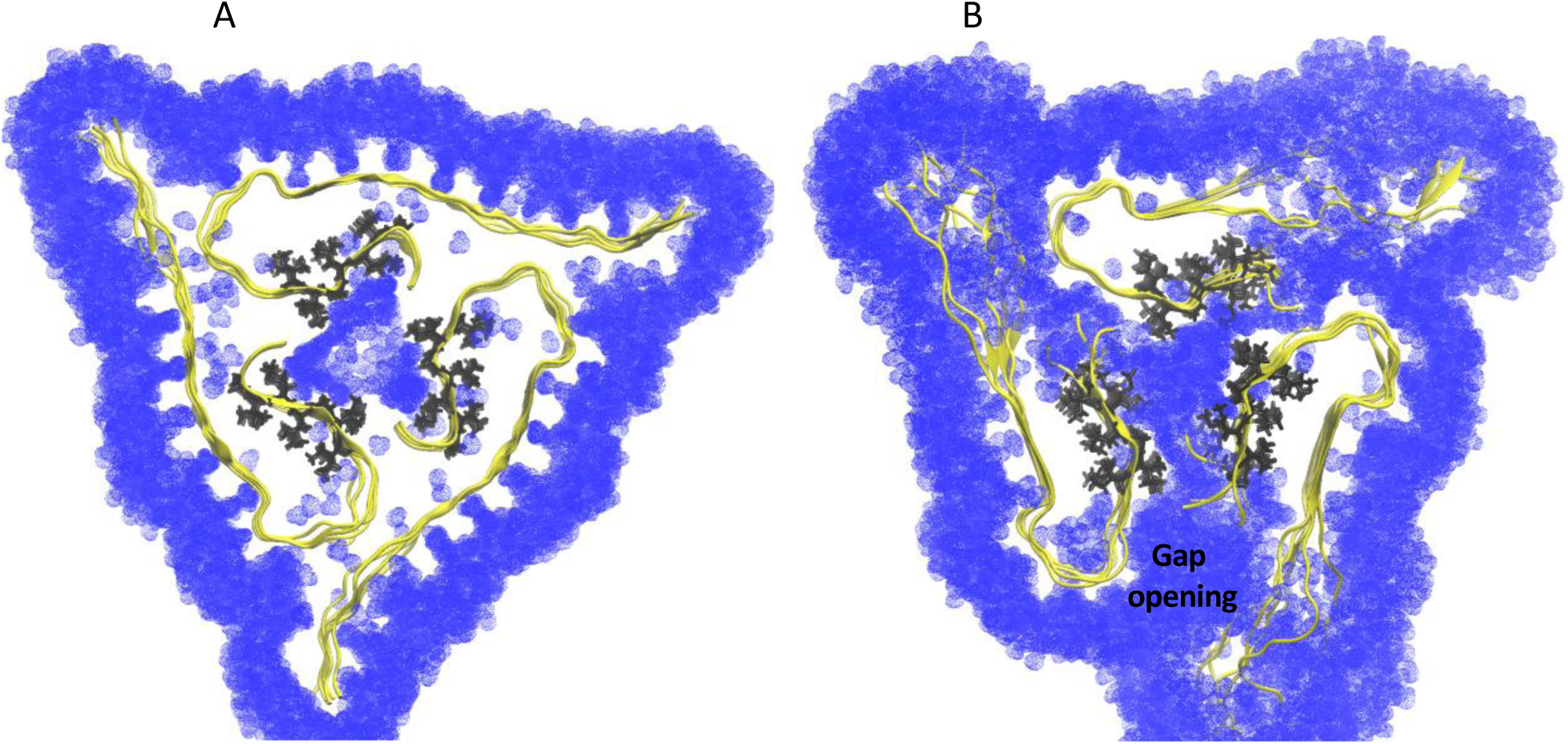
Water molecules inside and around the brain-seeded (2M4J) structure: initial (A), final (B). Only water molecules within 8 Å of the protein atoms are shown. Residues 31-37, some of which form the central core, are shown in black. Similar results were observed in the second replicate simulation of the 2M4J infinite system.

In the infinite-length 2M4J fibril, water was able to flow laterally through these gaps, which, in effect, formed an aqueous pore, with a continuous water phase extending into the central core of the fibril. The regions in which these gaps formed largely overlaps with the portion of the fibril previously shown to undergo relatively rapid hydrogen-deuterium (H/D) exchange in NMR experiments, i.e., A2, F4, D7, S8, G9, G25, and S26 (37). At these gaps, water was observed to flow in both the hydrogen bonding (*z*-axis, longitudinal) and side-chain (*xy*-plane, lateral) dimensions of the fibril. For the infinite fibrils, water initially enters the central core from the side-chain (radial) dimension (by construct, water cannot initially flow from bulk phase to the central core from the z-dimension because the fibril is infinite in this dimension). In contrast to the brain-seeded fibrils (2M4J), in the all-synthetic fibrils, this gap was much less apparent (2LMP trajectories) or was absent (2LMN simulations). The all-synthetic infinite fibril types had relatively water-tight edges. **Figure 4B** shows the flux of water molecules through the infinite-length 2M4J fibrils in the *x-y* plane. After an initial set of transient events in the region of residues 31-37 surrounding the central cavity, described below, new water molecules entered the core, while water molecules initially present gradually exited. The core radius was increased by ∼ 1.5Å by the end of the simulation reaching 11.0 Å (**Figure 5A,** in the longer second replicate simulation).

**Figure 5.**
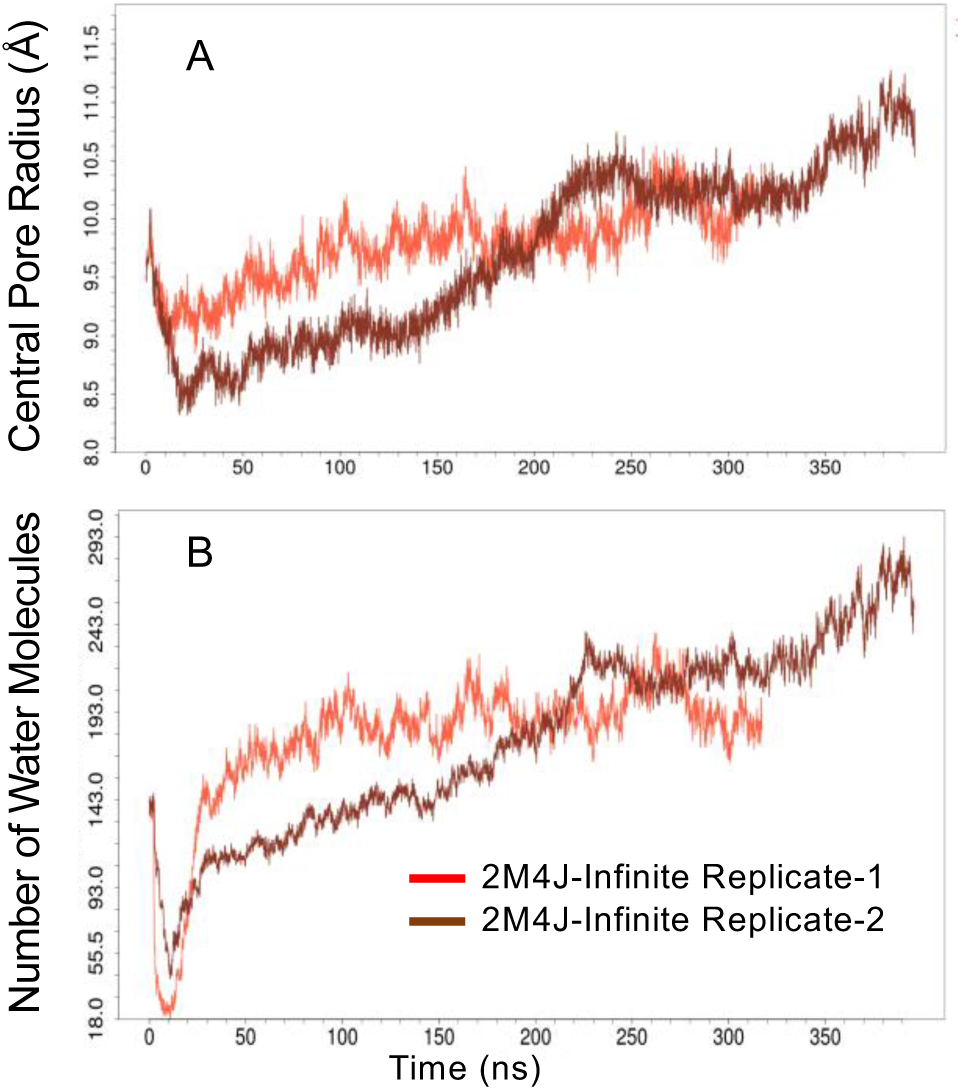
(A) The central core radius of the 2M4J structure in the course of the simulation (through the model-building, equilibration and production) and (B) the number of water molecules inside the central core. The reported radius was calculated as the average of radii corresponding to the middle three monomers that form the core for each frame of the trajectory.

**Figure 5B** shows the number of water molecules inside the central core of the 2M4J structure over the course of the simulation (through the model-building, equilibration and production phases). The central core was filled with water at the beginning of the simulation (**Fig 4A**). The number of water molecules decreased sharply but transiently at the beginning of the simulation, during the equilibration phase, and very quickly recovered its initial value. During this period, the protein backbone is restrained and gradually released to minimize the conformational changes while the side-chains equilibrated with water molecules. As the protein backbone restraints were released, these residues moved away from the center of the core causing a transient increase in the core radius (**Fig 5B**), and efflux of water molecules from the central core formed by residues 31-37. Water molecules refilled the core within ∼ 10 ns as the core structure adjusted to removal of the restraints. As the gaps opened at the sides of the fibrils and external water molecules entered the core, the initial water molecules gradually exited. By ∼ 100 ns, the number of water molecules in the 2M4J central core increased by ∼ 33% in replicate one (by ∼ 200 ns in the second replicate). It took 0.43 ± 0.12 ns (mean ± SD) for a water molecule to leave the central core of the 2M4J structure. Similar changes, although smaller, were observed for 2LMP infinite and finite fibrils. The central core of this system was also filled with water molecules, but its radius almost stayed constant throughout the simulation (Supporting Figure 6).

In order to compare diffusion through and within the 2M4J fibrils to diffusion in bulk water, the data were analyzed according to the equations for linear and anomalous diffusion processes (see Supporting Information for further details). **Figure 6** shows the mean square displacement (MSD) along the *x*, y, and *z* directions of water molecules beginning in the hydrated central core of the 2M4J infinite fibril as a function of lag time. ⟨*MSD*_*l*_⟩ is the ensemble average of the mean squared displacement of a given group of water molecules. In the *z* direction, ⟨*MSD*_*l*_⟩ for water within the 2M4J central core is linear with time, indicative of free diffusion, i.e., it followed the equation

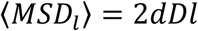

where *d* is the dimensionality of the translation (i.e., *d* = 3; 2; 1), *D* is the diffusion coefficient, and *l* is the lag time in the simulation. The diffusion coefficient for bulk water and water along the *z*-axis of the fibril were 3.96 × 10^-9^ and 3.73 × 10^-9^ m^2^/s, respectively. This is similar to that of TIP3P water model (4.0 × 10^-9^ m^2^/s) bulk solvent ^55^. For the *x* and *y* directions, water flux was slightly sub-diffusive, i.e., it followed the equation

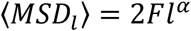

where F is a mobility factor (56), with the units of a diffusion constant. For *α* < 1, *α* = 1 or *α* > 1, the particles are sub-diffusive, regularly diffusive or super-diffusive, respectively. F was quite similar to D (4.38 × 10^-9^ and 4.48 × 10^-9^ for the *x* and *y* directions, respectively), and *α* was less than one (0.886 and 0.887 for the *x* and *y* directions), indicating slightly sub-diffusive flow. As expected, given the isotropy of the core, the parameters in *x* and *y* dimensions are similar. The sublinearity is attributable mainly to interactions water with amino acid side chains. In addition, it can be attributed to obstacles to translation in the *x-y* plane outside of the core, e.g., fibrils in adjacent images. These factors somewhat retard water diffusion along the *x* and *y* directions from matching that of bulk for longer lag times (**Figure 6**). Considering that the fibril is relatively permeant in the x-y plane due to the lateral pores, the slight sub-linearity is reasonable.

**Figure 6.**
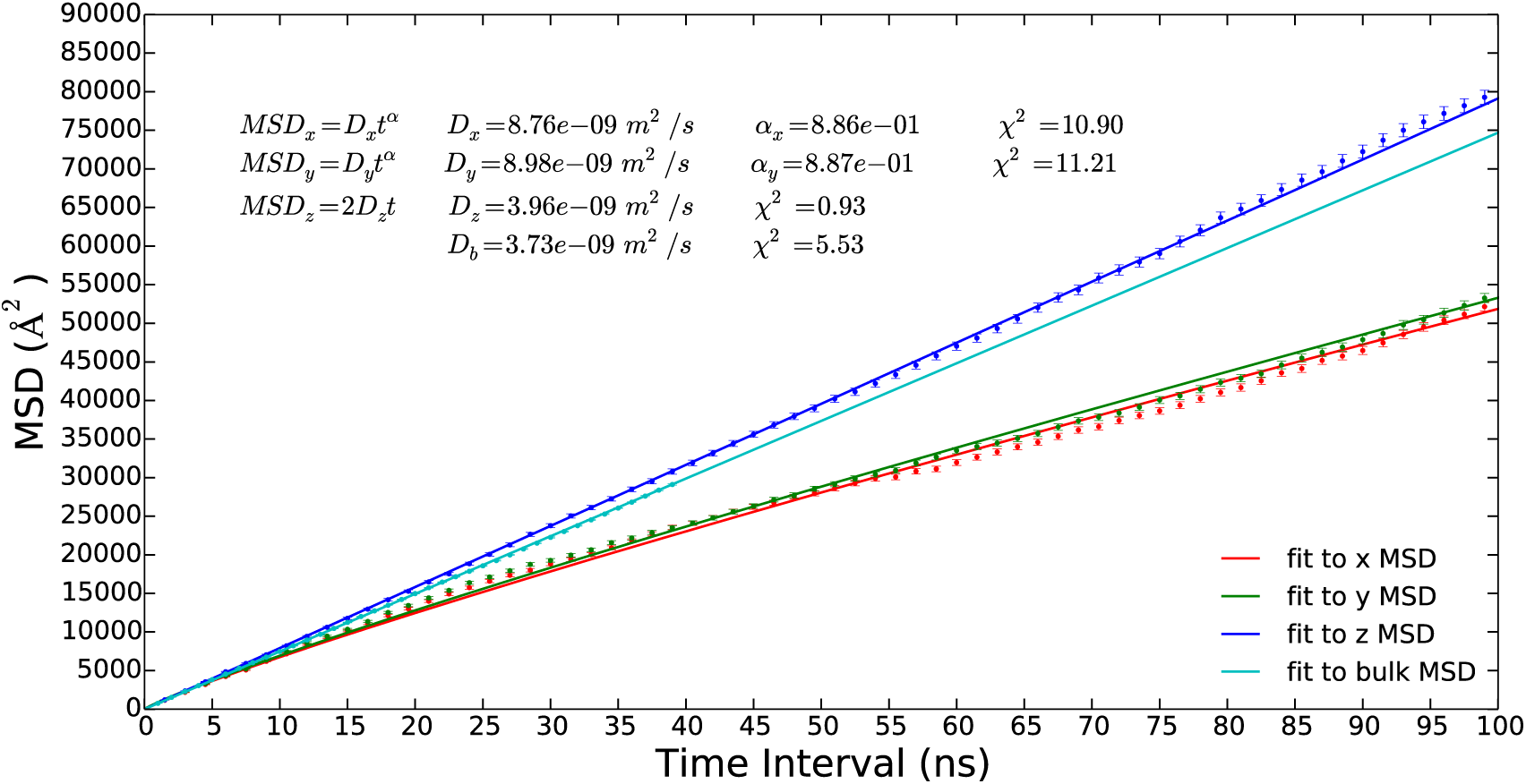
The mean square displacement (MSD) along the x, y and z directions for water molecules beginning in the hydrated central core of the 2M4J infinite fibril as a function of lag time, as well as that for an isotropic bulk water simulation; they are shown in red, green, blue and cyan respectively. For clarity, every hundredth point is plotted. The equations for linear and anomalous diffusion processes are given in Supporting Information.

The solvent contains only a small number of ions, added to maintain neutrality of the system. Nonetheless, we were able to track the passage of four sodium ions from the exterior of the 2M4J infinite fibrils (in the 500 ns long trajectory), into the fibril and the central core before exiting through a lateral core (**Supporting Figure 6** and included movie). Although this small number precludes quantitative measurements of ion diffusion rates, these observations suggest that ions could follow water into and through 2M4J fibrils.

### Secondary structure and hydrogen bonds

β-sheet content of all of structurers stayed relatively constant throughout the simulations (Supporting Figure 8A). Although finite fibrils deviated from the starting structures, they maintained a constant β-sheet content. Similarly, despite the gaps that opened at the sides of 2M4J fibrils during simulations, and the loss of symmetry, β-sheet stacks remained fairly constant. Similar pattern was observed in the number of backbone Hydrogen bonds (**Supporting Figure-8B).**

### Comparison of the simulation to the NMR data

**Supporting Figure 9** A shows histograms of the observed distances between atoms constrained in the NMR structure for both the NMR model, 2M4J, and the average of the last 200 time steps of the simulation in blue. The value of the constraint range is shown as a red box. The structure is considered to be consistent with the constraint if at least one distance is with the constraint range. By this criterion, we find that all the distance constraints are consistent with both the NMR structure while the simulated structure violates 4 constraints.

Supporting Figure 8 B shows the Ramachandran Plots for all backbone torsion angles constrained by the NMR chemical shift data. The chemical shift data constrain the torsion angles to the pink boxes in the figure, while the measured values from the NMR and simulated structures are shown by blue and black circles. If the majority of the measured values of the backbone torsion angles are consistent with the NMR constraint, the structure is considered to be consistent with the constraint. We find that 62.5% of the constraints are consistent with both the NMR and simulated structures, 12.5% are consistent with only the NMR structure, and 15.6% only with the simulated structure. 9.4% of the constraints are violated by both the NMR and simulated structures.

## Discussion

We compared three structural models of Aβ(1-40 residues) fibrils using MD simulations, two all-synthetic fibrils (2LMN, with a two-fold axis of symmetry; and 2LMP, with a three-fold axis of symmetry along the z-axis), and one brain-seeded fibril (2M4J, with a 3-fold axis of symmetry). The following are our four main findings.

First, we compared finite length (6 layered) fibrils to infinite fibrils constructed by replicating the structures along the *z*-axis using periodic images. In finite fibril simulations, fibril ends undergo more movement than interior regions, as indicated by a larger increase in RMSD and RMSF values – an effect not observed for infinitely long fibrils. The unraveling we observed for finite length fibrils may partly reflect the process of fibril disassembly from the fibril ends. The studies of infinitely long fibrils report more on the body of the mature fibril, i.e., without effects attributable to fibril ends.

Finite-length constructs may be useful as models of soluble oligomers or so-called “protofibrils” of Aβ(1-40 residues), i.e., precursors to the mature fibril. There is currently little structural information on soluble oligomers, but this limited information suggests structural similarity of some oligomers (57-60) to mature fibrils, i.e., parallel, in-register β-sheet. (This is not necessarily true of all oligomers, as other structures, i.e., anti-parallel β sheets, have also been reported (61-63).

Second, the D23-K28 salt bridges in initial structures of 2LMN and 2M4J generally were retained during the simulations. For 2LMP, where no salt bridges are initially present, either no or one salt bridge transiently formed during simulations. Distinguishing between intra- and intermolecular contacts by solid-state NMR requires experiments specifically designed to make such distinctions, e.g., isotope dilution experiments; such experiments are also of limited sensitivity. In our simulations, salt bridges in 2LMN and 2M4J (brain-seeded fibrils) were both intra- and/or intermolecular, i.e., K28 of one Aβ(1-40 residues) could interact simultaneously with D23 of the same or an adjacent Aβ(1-40 residues) molecule (along the *z*-direction). Thus, the number of salt bridges depended primarily on the initial conditions, indicating that these salt bridges are stable over the course of the simulations.

Third, molecules in each of the Aβ(1-40 residues) fibrils retain or develop a stagger in the *z*- direction. In 2LMN and 2LMP, this stagger is present in the initial structural model. In 2M4J fibrils, the N- and C-terminal β-strands of single Aβ(1-40 residues) molecules are initially coplanar. Over the course of the simulations, these fibrils tend to develop a stagger reminiscent of those in the other fibril types. This behavior was most evident in the finite length fibrils, but also occurred, to a limited extent, even in the infinitely long 2M4J fibrils. The inter-monomer salt bridge contributes to the stagger formation (Supporting Figure 2B). A consequence of the stagger is that the Aβ(1-40 residues) molecules at the ends of the fibrils have solvent-exposed hydrophobic amino acids side chains, as well as unsatisfied hydrogen bond donors and acceptors. As a result, the finite length fibrils exhibited much greater unraveling during the simulations than the infinitely long fibrils. Extrapolating to fibril assembly and disassembly, these results underscore the importance of the fibril ends in these processes. In addition, these results may help to rationalize the many observations that soluble oligomers and “protofibrils” are more cytotoxic than long fibrils: the fibril ends appear to be more mobile and reactive than the centers of fibrils.

Finally, and perhaps most importantly, the vertical stacks of the infinitely long 2M4J fibrils separate in a way that allows lateral entry of water into the fibril’s central cavity, and movement in the *z* direction (we observed the same phenomenon in simulations of 2LMP fibrils, though the changes were less marked). As expected, the β-sheet regions of the amyloids remain largely anhydrous and have essentially water-tight edges in all three structural forms. The 2M4J brain-seeded fibril, however, rapidly developed gaps between the N-terminus of one peptide molecule and the bend region of an adjacent peptide molecule. The residues comprising the gaps coincided with those that showed relatively rapid H/D exchange in SS-NMR studies of the fibrils, i.e., A2, F4, D7, S8, G9, G25, and S26.

The gaps observed in 2M4J allowed water to flow bidirectionally from the bulk phase, through the lateral gaps and into the central longitudinal core of the fibril. Recent studies using multidimensional IR have documented the presence of mobile water molecules in amyloid fibrils, even in relatively anhydrous areas (64, 65). Diffusion of water within the core in the *z* dimension of the fibril was similar to diffusion of bulk solvent, while slightly sub-linear diffusion was observed for movement in the *x-y* plane. Ions were also able to pass through the aqueous pores, and the few that permeated the core traveled *pari passu* with water, along the *z* dimension of the fibril (Supporting **Fig7**, movie). Alred et al. (66) used multi-layered 2lMP and 2M4J fibrils (not infinite) and did not observe similar gap openings. This may be due to their MD protocol where they scaled the mass of the solvent atoms by a factor of 0.5 (“reducing the viscosity of the solvent”, as stated in their paper). They also used a different version of Charmm forcefield. The new version (charmm36) has major improvements for the protein parameters.

Three points about SS-NMR can help to account for the fact that the structure of brain-seeded (2M4J) fibrils changed during simulation. First, SS-NMR (like solution NMR) examines events occurring on the microsecond or longer time scale, whereas our simulations examine events on the pico- and nanosecond time scale. Thus, some of the structural fluctuations observed in simulations could be averaged out or otherwise invisible in SS-NMR measurements. Second, the structural model proposed largely on the basis of SS-NMR data (37) includes a three-fold axis of rotational symmetry. This proposal of symmetry was based in part on the observation of a single spin system for each atom, which suggests a single conformation, or alternatively, static and/or dynamic inhomogeiety averaging to 3-fold symmetry. The presence of a single set of peaks is also true for SS-NMR data leading to the structural models of the all-synthetic fibrils, which show either a two- (2LMN) or three-fold (2LMP) axis of symmetry. Naturally, NMR cannot account for sparse variants that deviate from rotational symmetry, i.e., variants present at concentrations below the limits of detection. Thus, the proposal of rotational-symmetry is likely based on an averaging of detectable signals.

To determine the consistency of the MD simulations with the NMR structure of the brain-seeded fibrils (2M4J), we examined the NMR distance constraints torsional angle constraints and compared them to the major structural change of the fibril seen in the simulation. MD simulations show an opening of contacts between residues 5-10 and residues 24-27. The NMR constraint file has 10 contacts between these regions, but only one of them is unique; i.e. the other 9 can be assigned to contacts between other regions of the fibral that do not change much in the simulation. Thus, the only constraint violated by the simulated structure is between the N of residue 8 and the CG of residue 24. More experimental measurement will clear up this discrepancy.

The preceding findings on water flux through the fibril may have implications for the mechanisms of Aβ toxicity and AD pathogenesis in two areas. First, most investigators now believe that soluble Aβ oligomers and “protofibrils”, rather than mature fibrils are the more important neurotoxins in the pathogenesis of AD. Structural information on soluble Aβ oligomers is limited, however, and there is no consensus about what types of oligomers are most relevant to AD. As stated, both parallel, in-register and antiparallel β-sheet structures have been reported. Our simulations could shed some light on the former, which in their overall architecture resemble fibrils in that they contain a β-sheet – bend – β-sheet motif (22, 57, 58). Thus, some properties observed in the simulations for the fibrils (especially the finite length fibrils), may be applicable to the oligomers. In particular, the differences we observed between finite- and infinite-length fibrils highlight the differences between the structures of the fibril center and its ends: they illustrate the relative instability of the ends at which fibril dissolution probably occurs. Furthermore, they underscore the inherent instability of soluble toxic oligomers, which tend either to disassociate into monomers or grow into fibrils.

Second, one of the main proposed mechanisms of Aβ toxicity is the disruption of water and/or ion fluxes at neuronal cell membranes, especially of Ca^++^(67-76; for review see 77-80; for parallel references on islet amyloid polypeptide, see 81-85). While the precise mode of association of Aβ oligomers or fibrils with cell membranes remains poorly understood, it appears that Aβ peptides can insert into lipid bilayers, and otherwise disrupt cell membranes. Our simulations suggest that penetration of Aβ aggregates (fibrils or perhaps some oligomers) through cell membranes could open aberrant pores, where water and ions might flow *through* the fibril into or out of the cell. In addition to flow through the fibril, water might also flow through a defect in the membrane caused by the fibril or oligomer insertion. Flux through the fibril could represent an additional pathway for the abnormal ion transients that develop in the course of Aβ toxicity. This suggestion requires further experimental and modeling investigations.

## Conclusion

Our multiple ∼ 300 ns all-atom explicit solvent molecular dynamics simulations of three types of Aβ(1-40 residues) fibrils (2LMN, 2LMP, 2M4J) composed of either 6 or an infinite number of layers (made using periodic images) indicate that the former tend to unravel in a manner suggestive of the dissolution of fibrils or of some types of soluble oligomers. Salt-bridges in the structures tended to remain stable in those fibrils that contained them initially, while those without salt bridges did not develop them over the time course of the simulations. All fibrils showed some tendency to develop a “stagger” or register shift of β-strands along the fibril axis. Perhaps most importantly, the brain-seeded, 2M4J fibril rapidly develops gaps at its sides, which allows bidirectional flow of water and ions from the bulk phase in and out the central longitudinal core of the fibril. This behavior was also observed for the 2LMP fibrils, though to a lesser extent than for 2M4J fibrils. The development of an aberrant pore for water and ions could contribute to the neurotoxicity of Aβ aggregates. Since at least some oligomers resemble fibrils structurally, the previous statement could apply to soluble oligomers as well as fibrils.

## Author Contributions

E. J. Haddadian, S. C. Meredith, K. F. Freed and T. R. Sosnick designed the project. S. Natesh and E.J. Haddadian build the systems for simulation. S. Natesh ran the simulations and analyzed the data. J. R. Sachleben performed the NMR analysis. E. J. Haddadian and S. C. Meredith wrote the manuscript; all authors contributed to manuscript finalization.

## Acknowledgment

We like to thank Arthur Vale for running replicate simulations and data analysis. This work was completed using resources provided by the University of Chicago Research Computing Center, Computation Institute (in particular the Beagle and Midway computing cluster), Biological Sciences Division and Argonne National Laboratory under NIH grant S10 RR029030-01. This work was supported by NIH R01 AG048793 (SCM), the Alzheimer’s Association Zenith Fellowship Award (SCM), and NSF CHE-1363012 (KFF).

## Supporting figures

1. (A) All residue RMSD; finite fibrils are shown with open symbols, infinite fibrils with closed symbols. Red, blue and green correspond to 2LMN, 2LMP, and 2M4J, respectively. For clarity, only one of the replicates is shown and every fiftieth point is plotted. The RMSD for 2LMN, 2LMP, and 2M4J replicate simulations are shown is panels B, C, and D, respectively.

2. (A) Average number of D23-K28 salt bridges per stack. (B) Examples of intra- and inter-monomer salt bridge formations in infinite 2M4J structure are shown. The formation of inter-monomer salt brides can lead to stagger development. (C) The number of D23-K28 slat-bridges in the infinite fibril trajectory replicates.

3. The effect of 8 first residues in the stability of the structures. For fair comparison, the RMSD and RMSF values are calculated for residues 9-40 for the infinite-length 2LMP structure containing all 40 residues. The lack of these residues increased the structural deformation of the infinite-length fibril, evident from increases in the RMSD values in the infinite 2LMP system without the first 8 residues. The structural flexibility was also increased in one of the 32-residue fibril stacks (larger RMSF values in stack-1). In the RMSF plots, very flexible tail residues 37-40 are not shown. This region of the Aβ(1–40) contains a group of five charged residues that formed several transient salt bridges in the course of simulations helping to stabilize the 2LMP structure.

4. Stagger formation in the 2M4J infinite-length system. A stagger developed only in the third stack (refer to text **Figure 1** for the order of stacks). The C-terminal β-strand of peptide *i* that originally lay parallel to and coplanar with the N-terminal β-strand on the same peptide became coplanar with the N-terminal β-strand of peptide *i-1*.

5. (A) The central core radius of the 2LMP structure in the course of the simulation (through the model-building, equilibration and production) and (B) the number of water molecules inside the central core. The reported radius was calculated as the average of radii corresponding to the middle three monomers that form the core for each frame of the trajectory.

6. A sodium ion permeating through the infinitely long 2M4J structure in the course of the simulation. The starting position of the ion is shown in green and the final position in yellow. A movie of the sodium ion permeating through the infinitely long 2M4J structure is included. The ion begins outside of 2M4J structure and travels into and through the central core before exiting via a lateral pore. The unit cell contains a 6 layered 2M4J structure, and several periodic images along the fibril’s longitudinal axis are depicted.

7. (A) The percentile of β-sheets content for each residue in 2LMN, 2LMP and 2M4J simulations. (B) The number of backbone Hydrogen-bonds over simulation time for each system. The Hydrogen-bonds were identified with a distance (donor-acceptor) cutoff of 3.5 Å (or smaller) and an angle (donor-hydrogen-acceptor) cutoff of 35 degrees (or smaller).

8. Comparison of NMR structural constraints to NMR derived structure 2M4J and simulated structure. A) Histograms of interatomic distances for atoms constrained by NMR data.

Interatomic distances were measured between all intra and inter chain distances in the NMR and simulated structures. The counts for the histograms are shown in blue bars. The red box is the range of values consistent with the NMR distance measurement. B) Ramachandran Plot of the backbone torsion angles derived from the NMR and simulated structures. Pink boxes in the plots are the region of the torsion angle constrained by the NMR chemical shift data.

## References

(1) Alzheimer’s Association. Alzheimer’s Dement. 2015, 11, 332–384.

(2) World Health Organization. World atlas of ageing. Kobe, Japan: World Health Organization, Centre for Health Development, 1998.

(3) Plassman, B. L.; Langa, K. M.; Fisher, G. G.; Heeringa, S. G.; Weir, D. R.; Ofstedal, M. B.; Burke, J. R.; Hurd, M. D.; Potter, G. G.; Rodgers, W. L.; Steffens, D. C.; Willis, R. J.; Wallace, R. B. Neuroepidemiology Neuroepidemiology. 2007, 29, 125–132.

(4) Plassman, B. L.; Langa, K. M.; Fisher, G. G.; Heeringa, S. G.; Weir, D. R.; Ofstedal, M. B.; Burke, J. R.; Hurd. M. D.; Potter, G. G.; Rodgers, W. L.; Steffens, D. C.; McArdle, J. J.; Willis, R. J.; Wallace, R. B. Ann. Intern. Med. 2008, 148, 427–434.

(5) Hebert, L. E.; Scherr, P. A.; Bienias, J. L.; Bennett, D. A.; Evans, D. A. Arch. Neurol. 2003, 60, 1119–1122.

(6) Yan, R.; Fan, Q.; Zhou, J.; Vassar, R. Neurosci. Biobehav. Rev. 2016, 65, 326–340. (7)

(7) Hardy, J. J. Neurochem. 2009, 110, 1129–1134.

(8) Aguzzi, A.; O’Connor, T. Nat. Rev. Drug Discov. 2010, 9, 237–248.

(9) O’Brien, R. J.; Wong, P. C. Annu. Rev. Neurosci. 2011, 34, 185–204.

(10) Selkoe, D. J.; Hardy, J. EMBO Mol. Med. 2016, doi: 10.15252/emmm.201606210.

(11) Klein, W. L.; Krafft, G. A.; Finch, C. E. Trends Neurosci. 2001 24, 219–224.

(12) Caughey, B.; Lansbury, P. T. Annu. Rev. Neurosci. 2003, 26, 267–298.

(13) Klein, W. L.; Stine, W. B. Jr.; Teplow, D. B. Neurobiol. Aging. 2004, 25, 569–580.

(14) Teplow, D. B.; Lazo, N. D.; Bitan, G.; Bernstein, S.; Wyttenbach, T.; Bowers, M. T.; Baumketner, A.; Shea, J. E.; Urbanc, B.; Cruz, L.; Borreguero, J.; Stanley, H. E. Acc. Chem. Res. 2006, 39, 635–645.

(15) Glabe, C. C. Subcell. Biochem. 2005, 38, 167–177.

(16) Barghorn, S.; Nimmrich, V.; Striebinger, A.; Krantz, C.; Keller, P.; Janson, B.; Bahr, M.; Schmidt, M.; Bitner, R. S.; Harlan, J.; Barlow, E.; Ebert, U.; Hillen, H. J. Neurochem. 2005, 95, 834–847.

(17) Westermark, P.; Benson, M. D.; Buxbaum, J. N.; Cohen, A. S.; Frangione, B.; Ikeda, S.; Masters, C. L.; Merlini, G.; Saraiva, M. J.; Sipe, J. D. Amyloid 2007, 14, 179–183.

(18) Calabrese, B.; Shaked, G. M.; Tabarean, I. V.; Braga, J.; Koo, E. H.; Halpain, S. Mol. Cell. Neurosci. 2007, 35, 183–193.

(19) Martins, I. C.; Kuperstein, I.; Wilkinson, H.; Maes, E.; Vanbrabant, M.; Jonckheere, W.; Van Gelder, P.; Hartmann, D.; D’Hooge, R.; De Strooper, B.; Schymkowitz, J.; Rousseau, F. EMBO J. 2007. 27, 224–233.

(20) Irvine, G. B.; El-Agnaf, O. M.; Shankar, G. M.; Walsh, D. M. Mol. Med. 2008, 14, 451–464.

(21) Li, S.; Hong, S.; Shepardson, N. E.; Walsh, D. M.; Shankar, G. M.; Selkoe, D. Neuron 2009, 62, 788–801.

(22) Ahmed, M.; Davis, J.; Aucoin, D.; Sato, T.; Ahuja, S.; Aimoto, S.; Elliott, J. I.; Van Nostrand, W. E.; Smith, S. O. Nat. Struct. Mol. Biol. 2010, 17, 561–567.

(23) Funke, S. A. Int. J. Alzheimer’s Dis. 2011, 2011, 151645.

(24) Hayden, E. Y.; Teplow, D. B. Alzheimer’s Res. Ther. 2013 5:60, 1–11.

(25) Viola, K. L.; Klein, W. L. Acta Neuropathol. 2015 129, 183–206.

(26) Benzinger, T. L.; Gregory, D. M.; Burkoth, T. S.; Miller-Auer, H.; Lynn, D. G.; Botto, R. E.; Meredith, S. C. Proc. Natl. Acad. Sci. U. S. A. 1998, 95, 13407–13412.

(27) Tycko, R. Cold Spring Harb. Perspect. Med. 2016, 6, pii: a024083.

(28) Tycko R. Annu. Rev. Phys. Chem. 2011, 62, 279–299.

(29) Petkova AT, Leapman RD, Guo ZH, Yau WM, Mattson MP, Tycko R. Science. 2005, 307, 262–265.

(30) Paravastu, A. K.; Leapman, R. D.; Yau, W. M.; Tycko, R. Proc. Natl. Acad. Sci. U. S. A. 105, 18349–18354, 2008.

(31) Meinhardt, J.; Sachse, C.; Hortschansky, P.; Grigorieff, N.; Fändrich. M. J. Mol. Biol. 2009, 386, 869–877.

(32) Qiang, W.; Yau, W. M.; Tycko, R. J. Am. Chem. Soc. 2011, 133, 4018–4029.

(33) Kodali, R.; Williams, A. D.; Chemuru, S.; Wetzel, R. J. Mol. Biol. 2010, 401, 503–517.

(34) Colletier, J. P.; Laganowsky, A.; Landau, M.; Zhao, M.; Soriaga, A. B.; Goldschmidt, L.; Flot, D.; Cascio, D.; Sawaya, M. R.; Eisenberg, D. Proc. Natl. Acad. Sci. U. S. A. 2011, 108, 16938–16943.

(35) Paravastu, A. K.; Qahwash, I.; Leapman, R. D.; Meredith, S. C.; Tycko, R. Proc. Natl. Acad. Sci. U. S. A. 2009, 106, 7443–7448.

(36) Tycko, R. Protein Sci. 2014, 23, 1528–1539.

(37) Lu, J. X.; Qiang, W.; Yau, W. M.; Schwieters, C. D.; Meredith, S. C.; Tycko, R. Cell, 2013, 154, 1257–1268.

(38) Petkova, A. T.; Yau, W. M.; Tycko, R. Biochemistry, 2006, 45, 498–512.

(39) Petkova, A. T.; Leapman. R. D.; Yau W. M.; Tycko, R. Proc. Natl. Acad. Sci. U. S. A. 2008, 105, 18349–18354.

(40) Nasica-Labouze. J.; Nguyen, P. H.; Sterpone, F.; Berthoumieu, O.; Buchete, N. V.; Coté, S.; De Simone, A.; Doig, A. J.; Faller, P.; Garcia, A.; Laio, A.; Li, M. S.; Melchionna, S.; Mousseau, N.; Mu, Y.; Paravastu, A.; Pasquali, S.; Rosenman, D. J.; Strodel, B.; Tarus, B.; Viles, J. H.; Zhang, T.; Wang, C.; Derreumaux, P. Chem. Rev. 2015, 115, 3518–3563.

(41) Phillips, J. C.; Braun, R.; Wang, W.; Gumbart, J.; Tajkhorshid, E.; Villa, E.; Chipot, C.; Skeel, R. D.; Kalé, L.; Schulten, K. J. Comput. Chem. 2005, 26, 1781–1802.

(42) Eswar, N; Webb, B.; Marti-Renom, M. A.; Madhusudhan, S. M.; Eramian, M.; Shen, M. Y.; Pieper, U.; Sali, A. Current Protocols in Bioinformatics, 2006, Chapter 5:Unit 5.6.

(43) Humphrey, W.; Dalke, A.; Schulten, K. J. Molec. Graphics. 1996, 14, 33–38. (44)

(44) Hoover, W. G. Phys Rev A. 1985, 31, 1695–1697.

(45) Nosé S. J. Chem. Phys. 1984, 81, 511–519.

(46) Essmann U, Perera L, Berkowitz ML, Darden T, Lee H, Pederson L. J Chem Phys. 1995, 103, 8577–8593.

(47) Anderson, H.C. J. Comput. Phys. 1983, 52, 24–34.

(48) Smart, O. S.; Goodfellow, J. M.; Wallace, B. A. Biophys. J. 1993, 65, 2455–2460.

(49) Smart, O. S.; Neduvelil J. G.; Wang, X.; Wallace, B. A.; Sansom, M. S. P. HOLE: J.Mol.Graph. 1996, 14, 354–360.

(50) Sciarretta, K. L.; Gordon, D. J.; Petkova, A. T; Tycko, R.; Meredith, S. C. Biochemistry 2005, 44, 6003–6014.

(51) Nelson R.; Sawaya, M. R.; Balbirnie, M.; Madsen, A. Ø.; Riekel, C.; Grothe, R.; Eisenberg, D. Nature. 2005, 435, 773–778.

(52) Ivanova, M. I.; Sawaya, M. R.; Gingery, M.; Attinger, A.; Eisenberg, D. Proc. Natl. Acad. Sci. U. S. A. 2004, 101, 10584–10589.

(53) Nelson, R.; Eisenberg, D. Adv Protein Chem. 2006, 73, 235–282.

(54) Nelson, R.; Eisenberg, D. Curr. Opin. Struct. Biol. 2006, 16, 260–265.

(55) Price, D. J.; Brooks, C. L. III. J. Chem. Phys. 2004, 121, 10096.

(56) Demontis, P.; Stara, G.; Suffritti G. B. Microporous and Mesoporous Materials 2005, 86, 166–175.

(57) Chimon, S.; Ishii, Y. J. Am. Chem. Soc. 2005, 127, 13472–13473.

(58) Chimon, S.; Shaibat, M. A.; Jones, C. R.; Calero, D. C.; Aizezi, B.; Ishii, Y. Nat. Struct. Mol. Biol. 2007, 14, 1157–1164.

(59) Yu, L.; Edalji, R.; Harlan, J. E.; Holzman, T. F.; Lopez, A. P.; Labkovsky, B.; Hillen, H.; Barghorn, S.; Ebert, U.; Richardson, P. L.; Miesbauer, L.; Solomon, L.; Bartley, D.; Walter, K.; Johnson, R. W.; Hajduk, P. J.; Olejniczak, E. T. Biochemistry. 2009, 48, 1870–1877.

(60) Stroud, J. C.; Liu, C.; Teng, P. K.; Eisenberg, D. Proc. Natl. Acad. Sci. U. S. A. 2012, 109, 7717–7722.

(61) Gu, L.; Liu, C.; Guo, Z. J. Biol. Chem. 2013, 288, 18673–18683.

(62) Huang, D.; Zimmerman, M. I.; Martin, P. K.; Nix, A. J.; Rosenberry, T. L.; Paravastu, A. K. J. Mol. Biol. 2015, 427, 2319–2328.

(63) Gu, L.; Liu, C.; Stroud, J. C.; Ngo, S.; Jiang, L.; Guo, Z. J. Biol. Chem. 2014, 289, 27300– 27313.

(64) Kim, Y. S.; Liu, L.; Axelsen, P. H.; Hochstrasser, R. M. Proc. Natl. Acad. Sci. U. S. A. 2009, 106, 17751–17756.

(65) Ma, J.; Komatsu, H.; Kim, Y. S.; Liu, L.; Hochstrasser, R M.; Axelsen, P. H. ACS Chem. Neurosci. 2013, 4, 1236–1243.

(66) Alred E. J.; Phillips M., Workalemahu M. B.; Hansmann U. H. E.; Protein Sci. 2015, 24, 923–935

(67) Nagarathinam, A.; Höflinger, P.; Bühler, A.; Schäfer, C.; McGovern, G.; Jeffrey, M.; Staufenbiel, M.; Jucker, M.; Baumann, F. J. Neurosci. 2013, 33, 19284–19294.

(68) Hong, S.; Ostaszewski, B. L.; Yang, T.; O’Malley T. T.; Jin, M.; Yanagisawa, K.; Li, S.; Bartels, T.; Selkoe, D. J. Neuron. 2014, 82: 308–319.

(69) Tsai, H. H.; Lee, J. B.; Shih, Y. C.; Wan, L.; Shieh, F. K.; Chen, C. Y. ChemMedChem 2014, 9, 1002–1011.

(70) Yates, E. A.; Legleiter, J. Biochemistry. 2014, 53: 7038–7050.

(71) Qiang, W.; Akinlolu, R. D.; Nam, M.; Shu, N. Biochemistry. 2014, 53, 7503–7514.

(72) Ueno, H.; Yamaguchi, T.; Fukunaga, S.; Okada, Y.; Yano, Y.; Hoshino, M.; Matsuzaki, K. Biochemistry. 2014, 53, 7523–7530.

(73) Henry, S.; Vignaud, H.; Bobo, C.; Decossas, M.; Lambert, O.; Harte, E.; Alves, I. D.; Cullin, C.; Lecomte, S. Biomacromolecules. 2015, 16, 944–950.

(74) Malishev, R.; Nandi, S.; Kolusheva, S.; Levi-Kalisman, Y.; Klärner, F. G.; Schrader, T.; Bitan, G.; Jelinek, R. ACS Chem. Neurosci. 2015, 6, 1860–1869.

(75) Yi, X.; Zhang, Y.; Gong, M.; Yu, X.; Darabedian, N.; Zheng, J.; Zhou, F. Biochemistry. 2015, 54, 6323–6332.

(76) Peters, C.; Bascuñán, D.; Opazo, C.; Aguayo, L. G. J. Alzheimer’s Dis. 2016, 51, 689–699.

(77) Kotler, S. A.; Walsh, P.; Brender, J. R.; Ramamoorthy, A. Chem. Soc. Rev. 2014, 43, 6692–6700.

(78) Matsuzaki, K. Acc. Chem. Res. 2014, 47, 2397–2404.

(79) Andreasen, M.; Lorenzen, N.; Otzen, D. Biochim. Biophys. Acta 2015, 1848 1897–1907.

(80) Gorbenko, G.; Trusova, V.; Girych, M.; Adachi, E.; Mizuguchi, C.; Saito, H. Adv. Exp. Med. Biol. 2015, 855, 135–155.

(81) Cao, P.; Abedini, A.; Wang, H.; Tu, L. H.; Zhang, X.; Schmidt, A. M.; Raleigh, D P. Proc. Natl. Acad. Sci. U. S. A. 2013, 110, 19279–19284.

(82) Poojari, C.; Xiao, D.; Batista, V. S,; Strodel, B. Biophys. J. 2013, 105, 2323–2332.

(83) Kumar, S.; Birol, M.; Schlamadinger, D. E.; Wojcik, S. P.; Rhoades, E.; Miranker, A. D. Nat. Commun. 2016, 7, 11412.

(84) Junghans, A.; Watkins, E. B.; Majewski, J.; Miranker, A.; Stroe, I. Langmuir. 2016, 32, 4382–4391.

(85) Last, N. B.; Miranker, A. D. Proc. Natl. Acad. Sci. U. S. A. 2013, 110, 6382–6387.

